# FAIRshake: toolkit to evaluate the findability, accessibility, interoperability, and reusability of research digital resources

**DOI:** 10.1101/657676

**Authors:** Daniel J. B. Clarke, Lily Wang, Alex Jones, Megan L. Wojciechowicz, Denis Torre, Kathleen M. Jagodnik, Sherry L. Jenkins, Peter McQuilton, Zachary Flamholz, Moshe C. Silverstein, Brian M. Schilder, Kimberly Robasky, Claris Castillo, Ray Idaszak, Stanley C. Ahalt, Jason Williams, Stephan Schurer, Daniel J. Cooper, Ricardo de Miranda Azevedo, Juergen A. Klenk, Melissa A. Haendel, Jared Nedzel, Paul Avillach, Mary E. Shimoyama, Rayna M. Harris, Meredith Gamble, Rudy Poten, Amanda L. Charbonneau, Jennie Larkin, C. Titus Brown, Vivien R. Bonazzi, Michel J. Dumontier, Susanna-Assunta Sansone, Avi Ma’ayan

## Abstract

As more datasets, tools, workflows, APIs, and other digital resources are produced by the research community, it is becoming increasingly difficult to harmonize and organize these efforts for maximal synergistic integrated utilization. The Findable, Accessible, Interoperable, and Reusable (FAIR) guiding principles have prompted many stakeholders to consider strategies for tackling this challenge by making these digital resources follow common standards and best practices so that they can become more integrated and organized. Faced with the question of how to make digital resources more FAIR, it has become imperative to measure what it means to be FAIR. The diversity of resources, communities, and stakeholders have different goals and use cases and this makes assessment of FAIRness particularly challenging. To begin resolving this challenge, the FAIRshake toolkit was developed to enable the establishment of community-driven FAIR metrics and rubrics paired with manual, semi- and fully-automated FAIR assessment capabilities. The FAIRshake toolkit contains a database that lists registered digital resources, with their associated metrics, rubrics, and assessments. The FAIRshake toolkit also has a browser extension and a bookmarklet that enables viewing and submitting assessments from any website. The FAIR assessment results are visualized as an insignia that can be viewed on the FAIRshake website, or embedded within hosting websites. Using FAIRshake, a variety of bioinformatics tools, datasets listed on dbGaP, APIs registered in SmartAPI, workflows in Dockstore, and other biomedical digital resources were manually and automatically assessed for FAIRness. In each case, the assessments revealed room for improvement, which prompted enhancements that significantly upgraded FAIRness scores of several digital resources.

## Introduction

The Findable, Accessible, Interoperable, and Reusable (FAIR) guiding principles describe an urgent need to improve the infrastructure supporting scholarly data reuse, and outline several existing resources that already demonstrate various aspects of FAIR and associated driving technologies (1). A specific emphasis has been placed on ensuring that a machine could take advantage of adherence to the FAIR principles where the Resource Description Framework (RDF) (2) was suggested as the key globally-accepted framework for data and knowledge representation intended to be read and interpreted by machines. A key challenge in fulfilling the goals outlined by the FAIR guiding principles is the lack of consensus with respect to certain standards. In an effort to address some of these shortcomings, a comprehensive community-driven approach was taken to assemble FAIRsharing, a collection of standards, repositories, and policies (3). By collecting community-accepted elements of this kind, FAIRsharing can reveal domain-relevant community standards with respect to the FAIR principles. Filling important gaps, FAIRsharing has started to make it possible to realize the implementation of some FAIR principles by making standards FAIR. Several initiatives have already developed their own understandings of FAIRness and developed methods of assessing FAIRness by self- and peer-reviewed manual question-answer approaches (4,5). Because the different strategies for asserting FAIRness to date have been independent of one another, and as such not comparable, a template was created for constructing FAIR metrics around the original FAIR guiding principles (6). This FAIR metrics template is provided through GitHub in a format that can become community driven. The publication that describes the FAIR metrics contains self-evaluations by nine organizations. In the publication it was envisioned that a framework for automated evaluation of FAIRness could be devised using self-describing and programmatically executable metrics. The biomedical research community at large mostly embraces the FAIR guidelines; see, for example, a recent review by a group consisting of biopharma research and development representatives (7). While the FAIR metrics provide a concrete guide on how to begin to assess FAIRness, it is still unclear how to facilitate digital resource producers to define, assess, and implement FAIRness within their specialized specific projects. Here we present FAIRshake, a toolkit to systematically assess the FAIRness of any digital resource. The FAIRshake toolkit contains a database that enlists users, projects, digital resources, metrics, rubrics, and assessments. The FAIRshake toolkit also comes with a browser extension and a bookmarklet to enable viewing and submitting assessments from any website. The FAIR assessment results are visualized as an insignia that represent the FAIR score in a compact grid of squares colored in red, blue and purple. Below we briefly describe the various components of FAIRshake, and how the FAIRshake system has already been used to assess FAIRness for several projects.

## Results

### Overview

Overall, FAIRshake provides mechanisms to associate digital objects with rubrics and metrics to perform FAIR assessments. These assessments are communicated via the FAIR insignia (Fig. 1). FAIRshake also contains FAIR analytics modules that produce statistical reports about collections of assessments for a specific project (Fig. 2). The FAIRshake toolkit is composed of elements that include a full-stack web-server application containing a user interface, a backend database, and an application programming interface (API), as well as a Chrome extension and a Bookmarklet. The FAIRshake user interface is available from https://fairshake.cloud and the source code is openly available on GitHub (https://github.com/MaayanLab/FAIRshake). In an effort to make FAIRshake adhere to the FAIR guidelines, the FAIRshake endpoint-REST API are machine-readable with documentations for SmartAPI (8), Swagger/OpenAPI (https://swagger.io/), and CoreAPI (https://www.coreapi.org/). The API can be accessed via the human-friendly counterparts of these specifications with the REST Framework API explorer (https://www.django-rest-framework.org/topics/browsable-api/), Swagger UI (https://swagger.io/tools/swagger-ui/), and CoreAPI UI. A Jupyter Notebook (9) and YouTube tutorials (https://www.youtube.com/watch?v=7u0c4-yzXgA&list) are available to guide users through the process of using the FAIRshake interface and accessing FAIRshake programmatically.

**Fig. 1.**
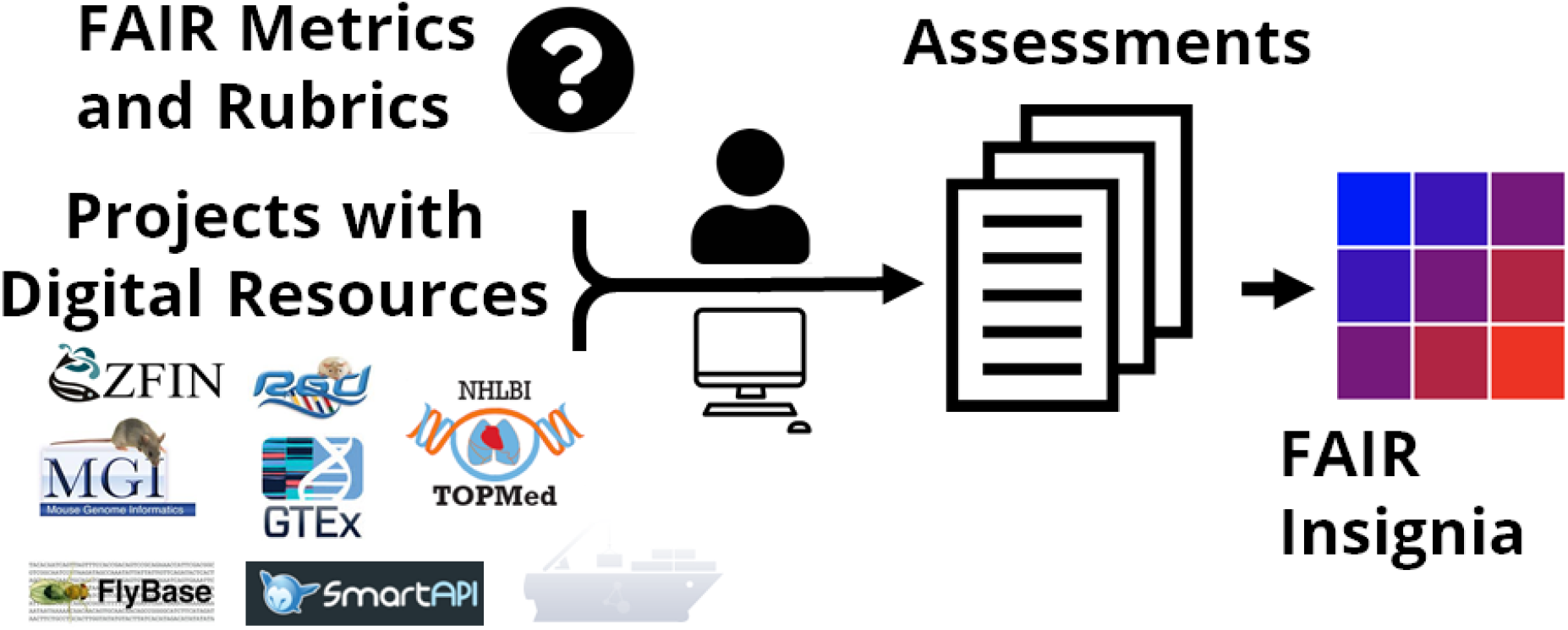
A diagram illustrating FAIRshake’s workflow. Digital resources from various projects are paired with FAIR metrics and rubrics to perform assessments that are visualized with the FAIR insignia.

**Fig. 2.**
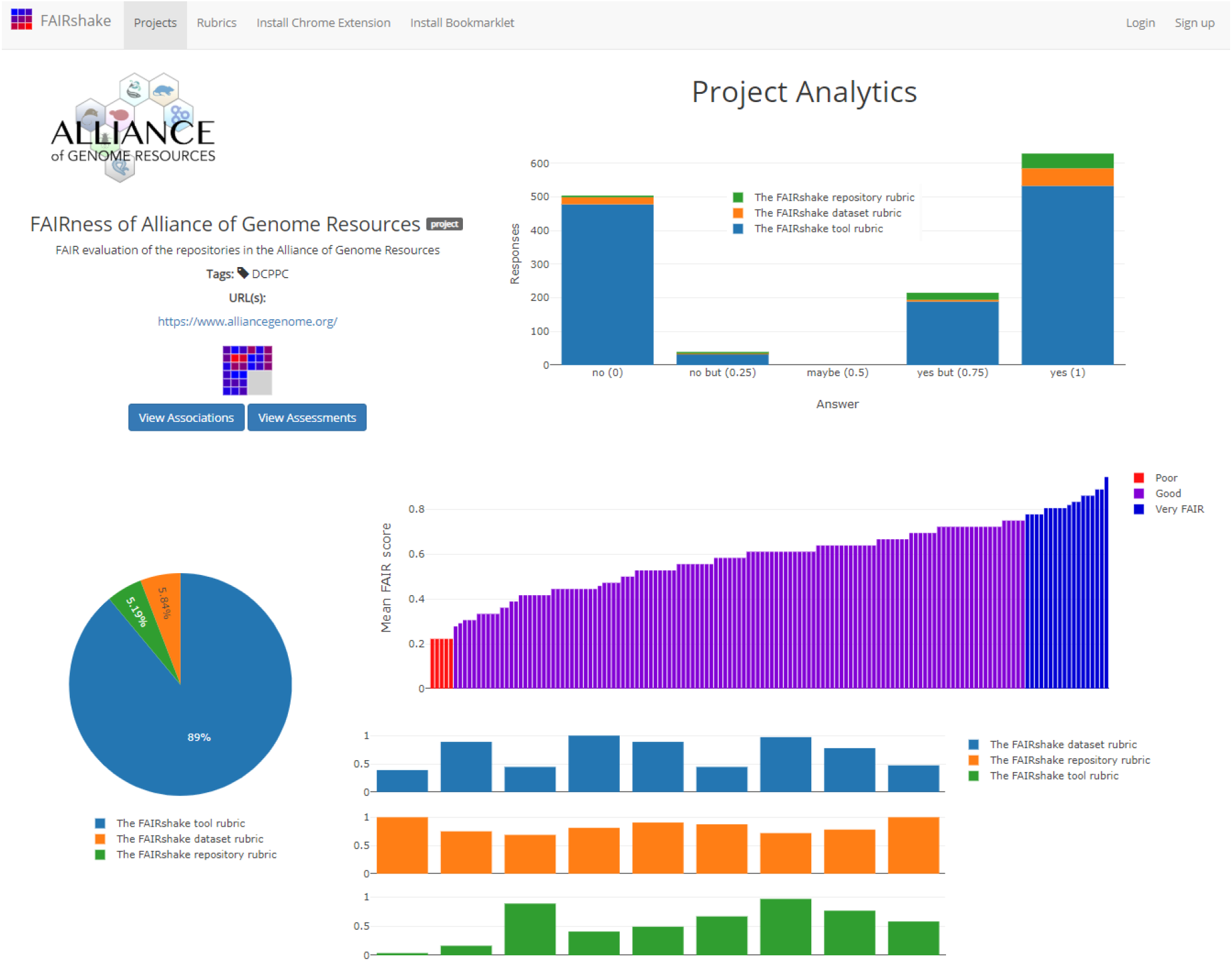
Each project within FAIRshake is provided with Project Analytics. These analytics include visualizations of the types of digital resources, the rubrics and metrics used to assess them, and the distribution of FAIR scores across all evaluated resources.

### Starting a project

To initiate a new project using FAIRshake, users are required to sign up. Sign up is available via standard registration and via OAuth (10) implementation of GitHub, ORCID (11), and Globus (12). Projects bundle a collection of thematically relevant digital resources. Project descriptions contain minimal information for identifying, displaying, and indexing the project. Within projects, users can assess the FAIRness of an arbitrary collection of digital resources. Project analytics are available to help a user better understand the overall FAIRness of the digital resources contained within the project. Project maintainers have access to all assessment values of the digital resources in their project. Projects enable users to group sets of digital resources and their assessments in a meaningful way. The same digital resources can have membership in any number of different projects. So far, FAIRshake contains 27 projects which are displayed as cards (Fig. 3).

**Fig. 3.**
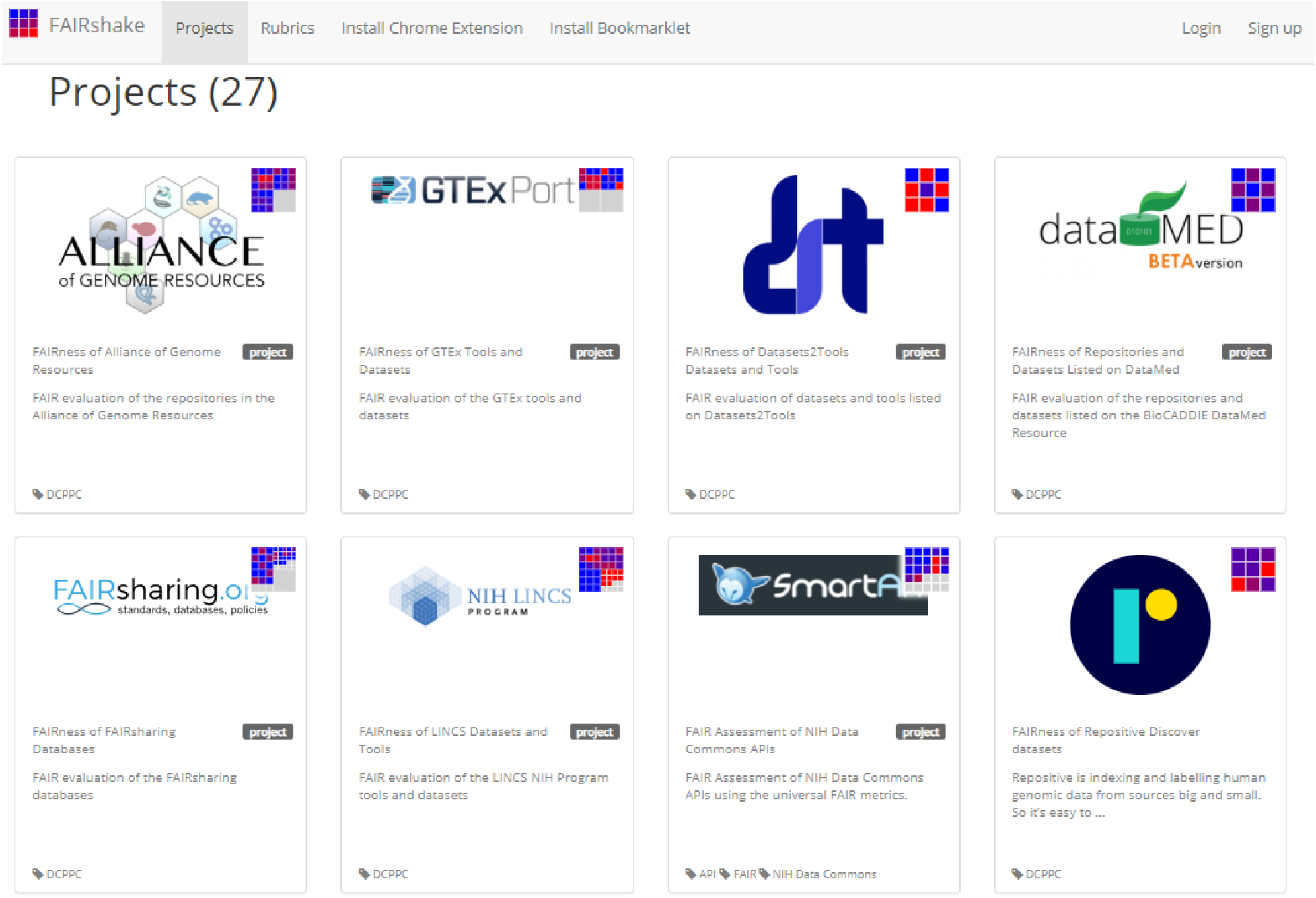
Screen capture from the projects tab of FAIRshake. FAIRshake displays projects as cards. Currently there are 27 projects within FAIRshake. Only 16 cards are shown.

### Registering digital resources

Each project in FAIRshake contains a collection of digital resources. FAIRshake handles all digital resources in the same way, regardless of whether they are datasets, tools, repositories, APIs, or any other type of digital resource. The minimum requirement for a digital resource to be registered with FAIRshake is one qualified URL identifying the digital resource. FAIRshake stores minimal optional information about each digital resource beyond the information needed for indexing, searching, and displaying the entry. The URL provided to describe each digital resource within FAIRshake is preferably a persistent URL that utilizes a community-accepted identifier system such as a Digital Object Identifier (DOI). This is important because the URL is the fundamental element for performing FAIR assessments.

### Metrics

FAIR metrics are questions that assess whether a digital object complies with a specific aspect of FAIR. A FAIR metric is directly related to one of the FAIR guiding principles. FAIRshake adopts the concept of a FAIR metric from the FAIRmetrics effort (6). In order to make FAIR metrics reusable, FAIRshake collects information about each metric, and when users attempt to associate a digital resource with metrics and rubrics, existing metrics are provided as a first choice. FAIR metrics represent a human-described concept that may or may not be automated; automation of such concepts can be done independently by linking actual source code to reference the persistent identifier of that metric’s semantic. Without linked code, metrics are simply questions that can be answered manually. FAIRshake defines several categorical answer types to FAIR metrics when manually assessed which are ultimately quantified to a value in a range between zero and one, or can take the property of being undefined. Programmatically, metric code can quantify the satisfaction of a given FAIR metric within this same range.

### Rubrics

The concept of a metric is supplemented with that of a FAIR rubric. A FAIR rubric is a collection of FAIR metrics. An assessment of a digital resource is performed using a specific rubric by obtaining answers to all of the metrics within the rubric. The use of a FAIR rubric makes it possible to establish a relevant and applicable group of metrics for a large number of digital resources, typically under the umbrella of a specific project. Linking rubrics to digital resources by association helps users to understand the context of the FAIR metrics that are best suited the digital resources in their projects. So far, 17 different rubrics are available via FAIRshake (Fig. 4).

**Fig. 4.**
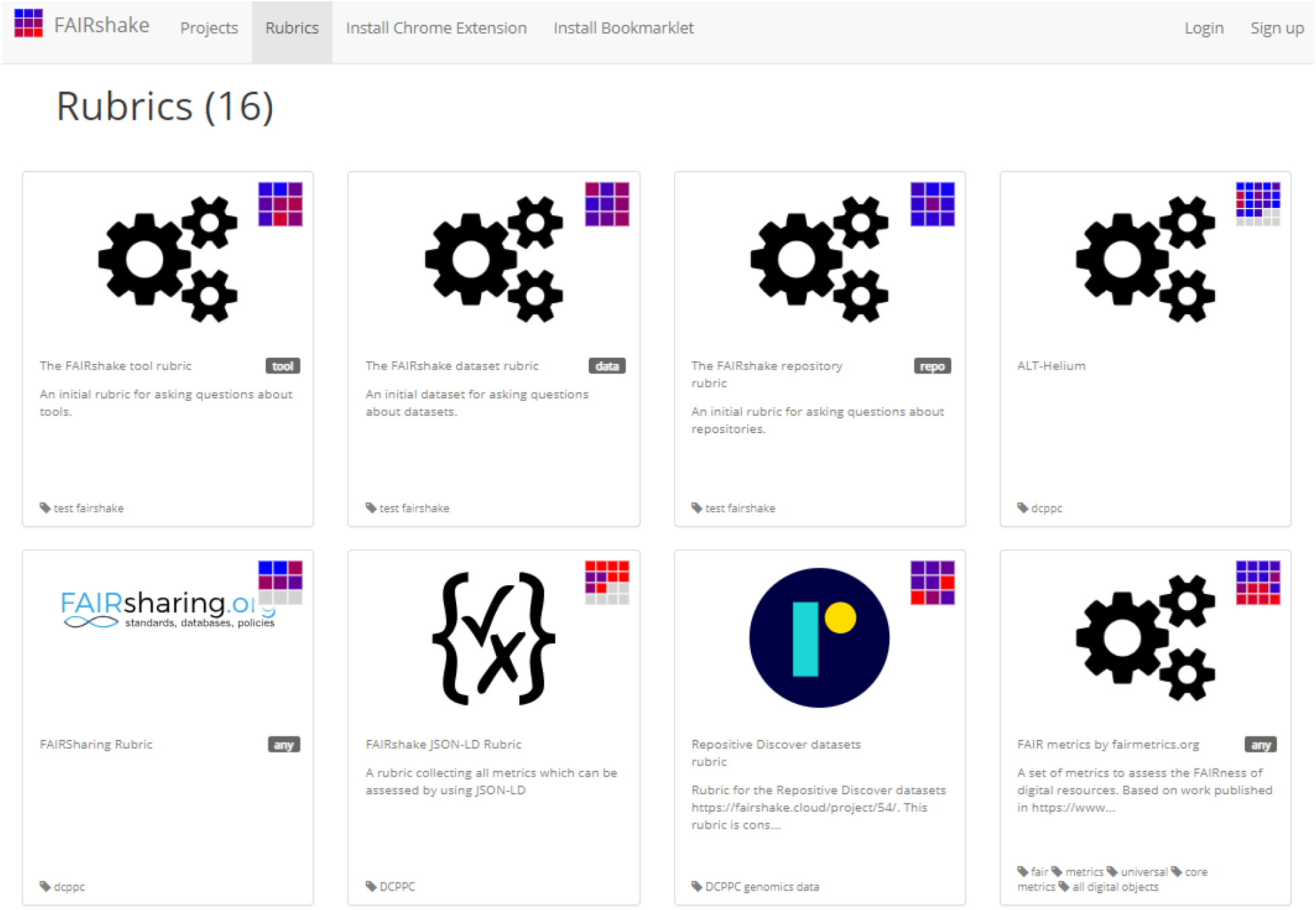
Screen capture from the Rubrics tab of FAIRshake. FAIRshake displays rubrics as cards. Currently there are 17 rubrics registered within FAIRshake.

### Assessments

An assessment can be performed on a digital resource that is coupled with a rubric. The assessment may involve the registration of a digital resource if it is not yet registered within FAIRshake. Metrics that have been codified can be automated as part of the assessment and will result in pre-populated fields when the user triggers the assessment manually. This is the case for semi-automated assessments, while automated assessments can only be triggered in bulk via the API. Leveraging on the usage of RDF, FAIRshake automatically extracts RDF-expressed schema.org metadata (13) from URLs with Extruct (14), a library for extracting embedded metadata from HTML markup. This approach is utilized by major search engines to index websites and bind information together. Using this RDF-expressed metadata alone, some FAIR metrics may be automatically resolved; including most of the exemplar universal metrics that were designed with RDF in mind (6). As schema.org expands its vocabulary through initiatives such as Bioschemas (15), RDF will enable these automatic mechanisms to scale FAIRshake assessments with the annotations present in public resources. Adopting other non-RDF based standards has also been accomplished with FAIRshake. In summary, any assessment of a digital resource within FAIRshake will attempt to obtain the answer to all metrics in the selected rubric, quantify those answers, and register the results in the FAIRshake database. The newly assessed digital resource will now have an associated insignia that reflects the results of the FAIR assessment.

### Insignias

The mechanism for visualizing FAIRness by FAIRshake is via the FAIRshake insignia (Fig. 1). The insignia uses a color gradient from blue (satisfactory) to red (unsatisfactory), visualizing how well a digital resource satisfied the FAIR metrics of the chosen rubric. Because the same digital resources can be assessed by different rubrics, composed of different metrics, the insignia dynamically expands to fit all assessments. If answers to the rubric are missing, the squares associated with these metrics will be colored in grey. Currently, the Insignia represents the mean of all associated assessments of a digital resource for each associated rubric. The mechanism for rendering the results of FAIR assessments, and visualizing digital resources’ state of FAIRness is packaged as a standalone JavaScript library for generating the insignia at any hosting website with few lines of code. The JavaScript library is available as a RequireJS package (https://requirejs.org/), or a nearly-pickled module (NPM) package (https://www.npmjs.com/), and code snippets are provided to demonstrate how it can be used. Through this library, a Chrome browser extension and bookmarklet were developed for rendering the visualization of FAIR insignias on any website without the need of the hosting site to modify their website’s source code.

### Case study 1

The first use case of FAIRshake involved the manual assessment of 376 bioinformatics tools by 23 participants. This evaluation was performed using a rubric composed of 9 questions/metrics (https://fairshake.cloud/rubric/7). These FAIR metrics capture various aspects of FAIRness that could be established by primarily examining the tool’s website. Of these tools, 132 of them were developed by members of the Alliance of Genome Resources (AGR) (https://www.alliancegenome.org/). Detailed results and breakdown of these assessments were captured in an HTML table and associated Jupyter Notebooks that are available at https://maayanlab.github.io/AGR-FAIR-Website/. Overall, we observed that the examined AGR tools scored well in regards to providing tools and data for download, use of ontologies, and providing contact information, while most AGR tools do not provide the source code, versioning information, or API access (Fig. 5).

**Fig. 5.**
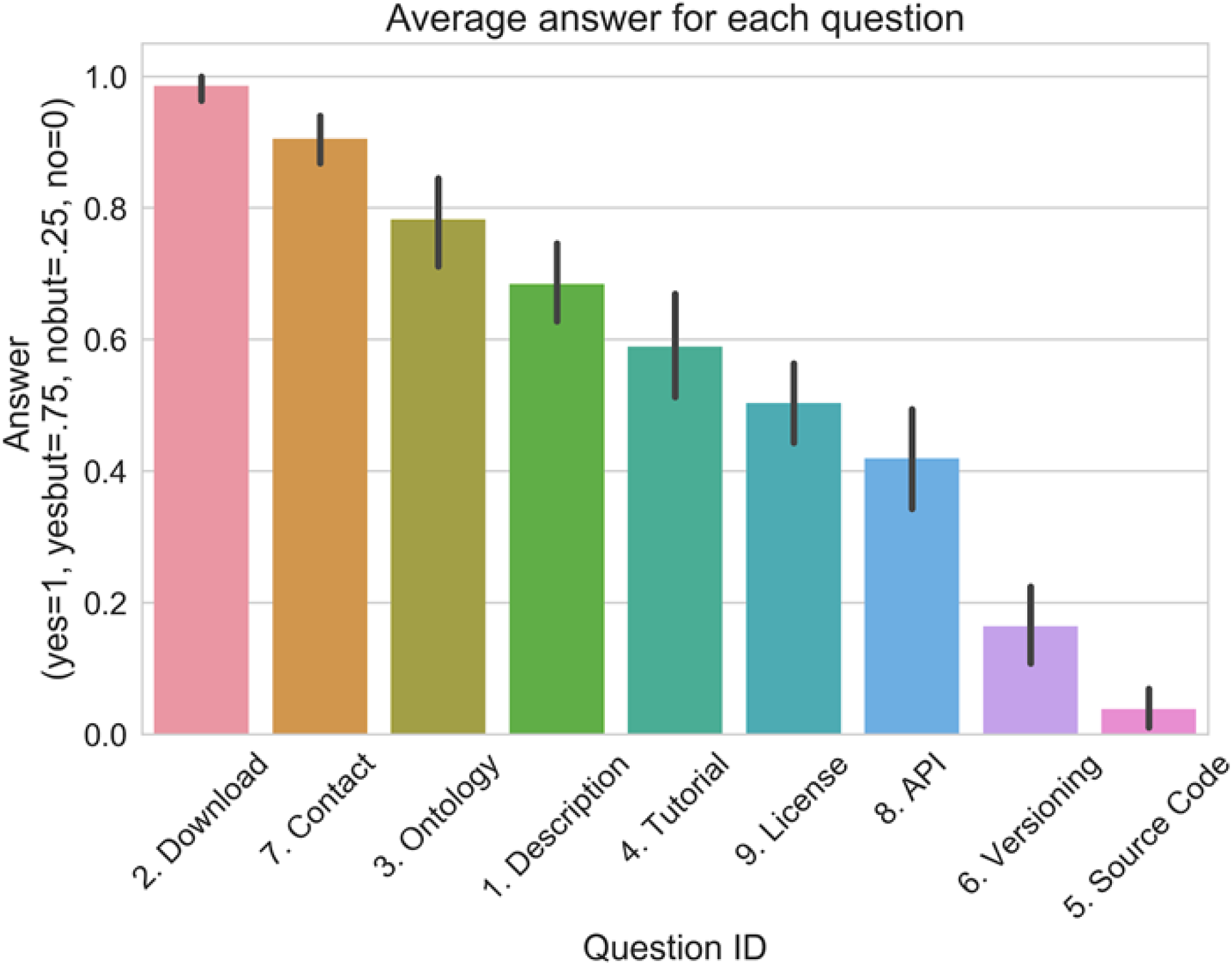
Distribution of average FAIR scores for 132 AGR tools assessed with an initial set of 9 FAIR metrics.

### Case study 2

To maximally utilize the efforts of the FAIRsharing team and minimize duplication, FAIRshake is integrated with the FAIRsharing [3] API for FAIR knowledge resolution. This is helping to drive semi-automated assessments relating to resources already available within FAIRsharing. Through this integration, FAIRshake has indexed a total of 1175 FAIRsharing repositories enabling user-driven manual and semi-automated assessments of those resources.

### Case study 3

Metadata related to the National Heart, Lung, and Blood Institute (NHLBI) Trans-Omics for Precision Medicine (TOPMed) studies are stored in the database of Genotypes and Phenotypes (dbGaP). Although the data in dbGaP is protected for privacy preservation, the metadata associated with the dbGaP database is available via a public open FTP site (ftp://ftp.ncbi.nlm.nih.gov/dbgap/studies/). This metadata follows a standardized directory structure capable of being codified. Paired with the use of the W3C standard XML Schema XSD; FAIRshake was able to automatically identify fields in the metadata that pertain to several FAIR metrics. Using these mappings, FAIRshake was employed to automatically assess the TOPMed studies in dbGaP using the information provided by the metadata. Although incomplete, this automated assessment revealed aspects of the metadata missing important information for FAIRness despite the presence of machine-readable metadata (Fig. 6).

**Fig. 6.**
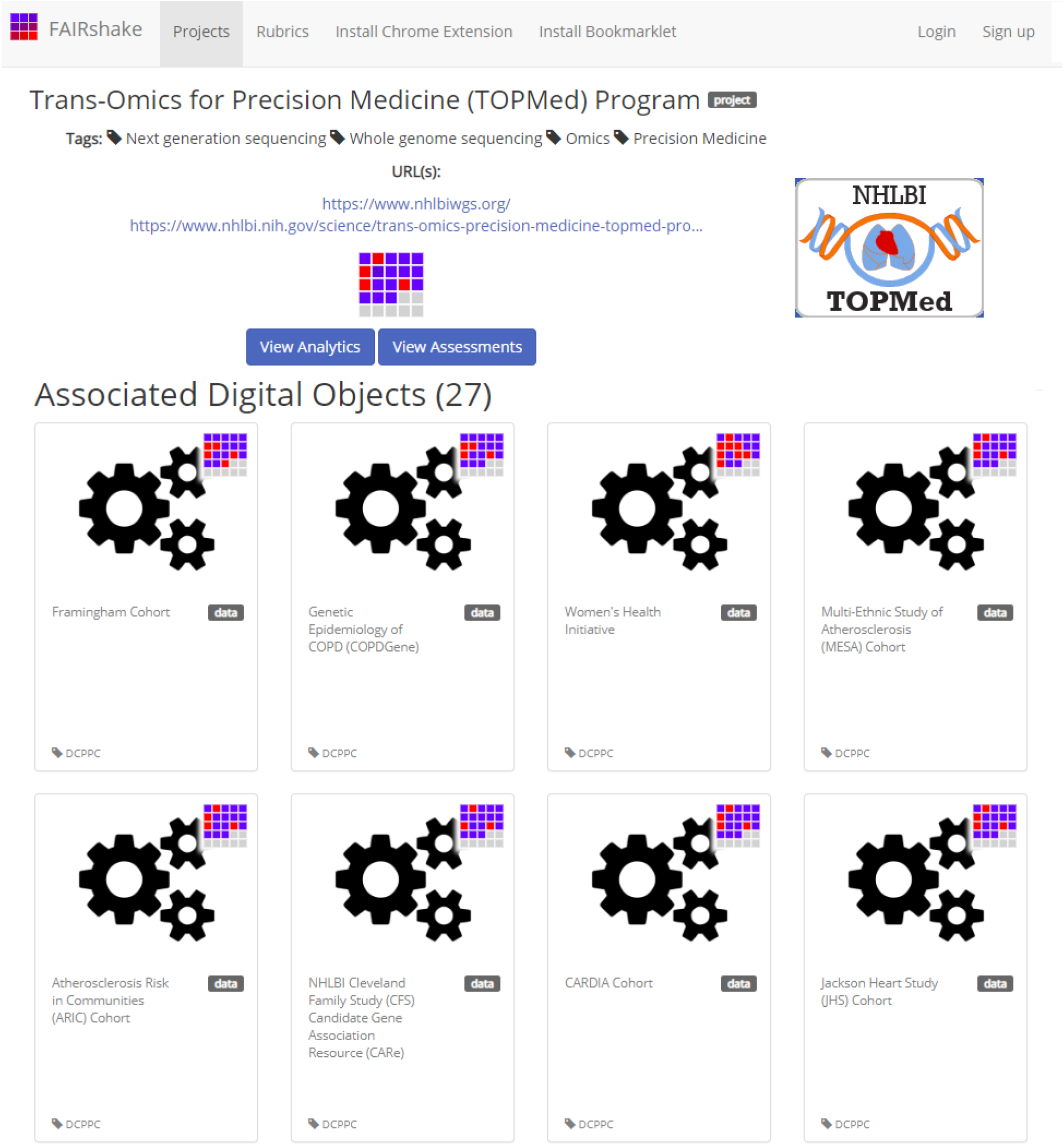
Screen capture from the automated FAIR assessments performed by FAIRshake to evaluate the FAIRness of TOPMed dataset entries in dbGaP.

### Case study 4

The SmartAPI resource maintains a repository of APIs with a machine-readable and validatable JSON-based documentation [9]. The machine-readable nature of this documentation structure enabled automatic resolution of attributes available, or not, within the structure pertaining to relevant FAIR metrics including aspects such as contact information, license availability, API best-practices, FAIR vocabulary utilization, and the presence of a Terms of Service statement. Altogether, 82 resources are registered with SmartAPI, and all were assessed for FAIRness by FAIRshake. Some of those resources have since improved their documentation, or their API, to improve their FAIR scores.

### Case study 5

As part of two 2-day NCBI-organized hackathons, participants used FAIRshake to evaluate the FAIRness of several NIH-related and other resources. The first hackathon took place on the NIH campus in Bethesda, MD in February 2019. The assessments from this hackathon are summarized assessments hosted on GitHub (https://github.com/NCBI-Hackathons/FAIRy-Compass). Relevant metrics were gathered by participants and 227 repositories were automatically registered and assessed using FAIRshake. Automated assessments were made possible by taking advantage of GitHub’s rich API. Approximately half of the metrics that were identified could not be automatically assessed with the available machine-readable information and, due to time constraints, were deferred to be established at a later time. The second hackathon took place as part of the BioIT World annual meeting in Boston, MA in April, 2019. Teams registered their projects and digital resources in FAIRshake and evaluated them before and after the hackathon. An example for such before and after assessment of the digital resource Exomiser (16) is provided (Fig. 7).

**Fig. 7.**
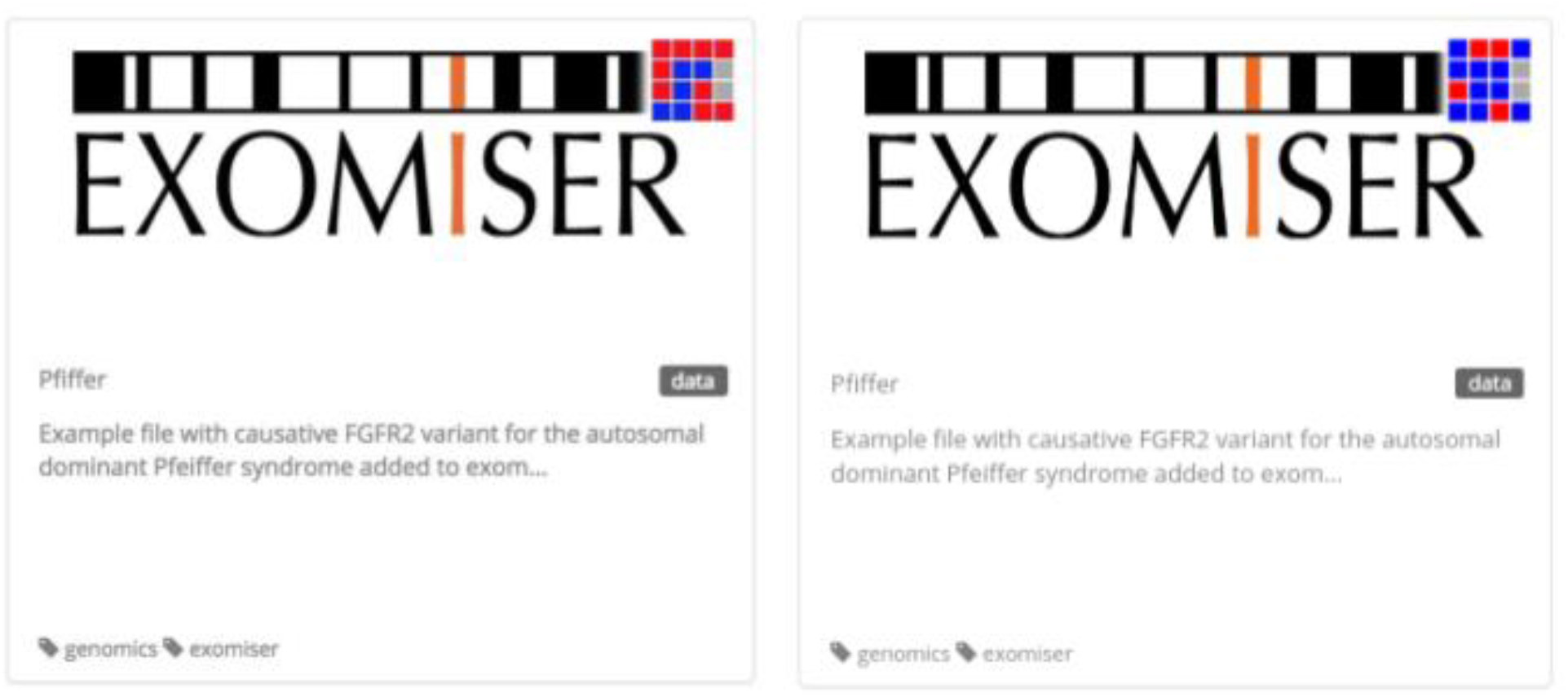
An example of a FAIRshake entry for EXOMISER, a digital resource that was assessed before and after the 2019 BioIT World NCBI FAIR Hackathon.

## Discussion

FAIRshake was developed as a toolkit to promote the FAIRification of digital objects produced by research projects. FAIRshake is not intended to judge or punish digital resource producers but rather to promote their awareness about standards and guide digital object producers to implement community-based best practices for their own benefit of attracting, retaining, and enabling more users. There is common confusion between assessing the quality of a resource and assessing its FAIRness. It should be made clear that FAIRshake was designed to assess FAIRness and a low FAIR score does not mean that a digital resource is lacking quality, usefulness, user-friendliness, or innovativeness. Another aspect of confusion about FAIR is the mistaken association of FAIRness with openness. Being FAIR does not entail making data, source code, tools, or any other digital resource, free and openly available. Rather, the FAIR guidelines only require that access and usage policies are provided and stated clearly (17,18).

By facilitating the creation of both manual and automated FAIR assessments, and enabling FAIR metric findability, FAIRshake promotes the involvement of more stakeholders. Starting with the process of manual FAIR assessments, the capacity for automation is expected to further expand as more adoption is realized. The findability of FAIR metrics within FAIRshake makes it possible to design community-adopted metrics that can be tested. FAIRshake strives to evolve with the community, adding new features to accommodate community demands as they arise while facilitating more assessments. With the approach of enabling the development of FAIR metrics and rubrics, the assessment of resources can happen in parallel; the community can start assessing FAIRness while still evolving the definition of what it means to be FAIR. FAIRshake facilitates dynamic metric re-use and provides analytical tools to understand the global and relative performance of resources and metrics. With transparency, FAIRshake enables the community to study the FAIRness of their resources they produce and use.

FAIRshake was developed to meet the demands of the biomedical research community. With integration of a number of community-accepted standards including RDF, FAIRSharing, SmartAPI, and schema.org, FAIRshake is already a versatile and capable toolkit for facilitating FAIR assessments on a diverse set of digital objects including data, tools, repositories, and APIs. Initial assessments have included 376 manual assessments of bioinformatics tools and databases, 27 automated assessments of NLBI TOPMed studies, 82 automated assessments of APIs registered on SmartAPI, 104 automated assessments of Dockstore resources, and 227 semi-automated assessments of NIH-related tools. These assessments resulted in improvements to FAIRshake’s capacity to perform assessments, enhancements to existing metrics for evaluating FAIRness, and upgrades to the resources that were assessed through an effort that was initiated based on the feedback received from the FAIRshake assessments. Throughout our initial assessments, it has become clear that many established community standards are not being employed within the biomedical research community, largely due to a lack of awareness of such standards. As the community continues to evolve towards better defining FAIRness, the FAIR metrics are expected to converge, and the FAIR assessments are likely to become more automated.

FAIRshake will continue to evolve with community demand. Continued improvements to the clarity, usability, and FAIRness of FAIRshake are planned. Similarly, integration with existing FAIR resources, such as FAIRSharing, will enable the display of assessments collected by FAIRshake on digital resource landing pages so that a broader community will become more aware of FAIRshake. The FAIRshake platform codebase can be reused for the assessment of other digital and physical products such as publications, events, books, and courses. However, such assessments may not be relevant to the FAIR guiding principles. Nevertheless, the FAIRshake platform is flexible enough that it can facilitate other related applications. One such potential future application is repurposing the FAIRshake codebase as a general platform for scientific peer review.

## Availability

The primary interface to FAIRshake is at: https://fairshake.cloud

The FAIRshake Chrome extension is available from: https://chrome.google.com/webstore/detail/fairshake/pihohcecpiomegpagadljmdifpbkhnjn?hl=en-US

The FAIRshake source code is available from GitHub at: https://github.com/MaayanLab/FAIRshake

## Acknowledgements

This work was partially supported by NIH grants OT3-OD025467 and U54-HL127624.

